# Phenotypic Age: a novel signature of mortality and morbidity risk

**DOI:** 10.1101/363291

**Authors:** Zuyun Liu, Pei-Lun Kuo, Steve Horvath, Eileen Crimmins, Luigi Ferrucci, Morgan Levine

## Abstract

**Background:** A person’s rate of aging has important implications for his/her risk of death and disease, thus, quantifying aging using observable characteristics has important applications for clinical, basic, and observational research. We aimed to validate a novel aging measure, “Phenotypic Age”, constructed based on routine clinical chemistry measures, by assessing its applicability for differentiating risk for morbidity and mortality in both healthy and unhealthy populations of various ages.

**Methods:** A nationally representative US sample, NHANES III, was used to derive “Phenotypic Age” based on a linear combination of chronological age and nine multi-system clinical chemistry measures, selected via cox proportional elastic net. Mortality predictions were validated using an independent sample (NHANES IV), consisting of 11,432 participants, for whom we observed a total of 871 deaths, ascertained over 12.6 year of follow-up. Proportional hazard models and ROC curves were used to evaluate predictions.

**Results:** Phenotypic Age was significantly associated with all-cause mortality and cause-specific mortality. These results were robust to age and sex stratification, and remained even when excluding short-term mortality. Similarly, Phenotypic Age was associated with mortality among seemingly “healthy” participants—defined as those who were disease-free and had normal BMI at baseline—as well as the oldest-old (aged 85+)—a group with high disease burden.

**Conclusions:** Phenotypic Age is a reliable predictor of all-cause and cause-specific mortality in multiple subgroups of the population. Risk stratification by this composite measure is far superior to that of the individual measures that go into it, as well as traditional measures of health. It is able to differentiate individuals who appear healthy, who may have otherwise been missed using traditional health assessments. Further, it can differentiate risk among persons with shared disease burden. Overall, this easily measured metric may be useful in the clinical setting and facilitate secondary and tertiary prevention strategies.

## Introduction

Rapid population aging represents a major public health burden, as aging is one of the leading risk factors for most chronic diseases [1, 2]. As a result, preventive strategies and interventions that promote healthy aging are critical. While everyone ages, that rate at which aging occurs is heterogeneous, and between-person variations in the pace of aging manifest as differences in susceptibility to death and disease. Thus, quantifying aging in individuals, particularly at younger ages, will facilitate secondary and tertiary prevention through earlier identification of high risk individuals or groups. However, a key issue remains in how to measure aging. Further, to be applicable to clinical setting, such assessments should be easily measurable using existing instruments; must do a better job at capturing risk stratification than current tools; and should be able to differentiate risk early, prior to manifestation of disease or disability.

One method for determining whether a person is younger or older than expected, based on his/her chronological age, is to compare observable characteristics—such as clinically observable data—to that of the general population. These characteristics, or “phenotypes”, are presumed to be manifestations of the biological aging processes that occur at the cellular and intracellular levels. A number of aging measures have been proposed using characteristics based on molecular measures [3, 4] and/or clinical chemistry measures [5–7]. Overall, the molecular measures exhibit weak associations with aging outcomes [8–14], compared to the clinical measures [5–7, 15–17], and are not currently feasible for use in the clinical setting. In contrast, the better performance and relative affordability and practicality of composite scores based on traditional clinical chemistry measures, may make them more suitable for evaluating effects of aging interventions or identifying groups at higher risk of death and disease.

Unfortunately, most of the existing aging biomarkers were generated based on associations between composite variables and chronological age—with no integration of information on how the variables influence morbidity and mortality. Further, given that individuals vary in their rate of aging, chronological time is an imperfect proxy for building an aging measure [18]. Recently, we developed a new metric, “Phenotypic Age”, that incorporates composite clinical chemistry biomarkers based on parametrization from a Gompertz mortality model. Rather than predicting chronological age—as previous measures have done—this measure is optimized to differentiate mortality risk among persons of the same chronological age, using data from a variety of multisystem clinical chemistry measures. In general, a person’s Phenotypic Age signifies the age within the general population that corresponds with that person’s mortality risk—two individuals may be 50 years old chronologically, but one may have the mortality risk of someone who is 55 years old, while the other may have the mortality risk of someone who is 45 years old.

The goal of this study was to validate this measure by: 1) assessing whether it is a robust predictor of all-cause mortality, compared to traditional risk factors; 2) establishing how it relates to various causes of death, and/or comorbid conditions; and 3) determining whether this new measure is predictive of long-term mortality risk in a healthy (i.e., normal body mass index [BMI] and disease-free) population, as well as a highly disease prevalent sub-sample of the population (i.e., the oldest-old).

## Methods

### Study population

We developed “Phenotypic Age” in a sample (n=9,926) from the third National Health and Nutrition Examination Survey (NHANES III) (1988-1994), a nationally representative cross-sectional study conducted by the National Center for Health Statistics. The subsequent validation sample used here was from the NHANES IV (1999-2010), which consists of independent cross-sections of the U.S. population taken every two years. Data for NHANES III and IV were collected from at-home interviews and examinations performed at a Mobile Examination Center (MEC). Details of recruitment, procedures, population characteristics, and study design are provided through the Centers for Disease Control and Prevention (CDC) [19]. Our validation sample (Table S1) included n=11,432 of the 12,640 NHANES IV participants (aged 20–84 y). Excluded participants are those with missing data on biomarkers or who did not complete at least eight hours of fasting. Those who were ages 85+ were also excluded from all analyses except age-stratified mortality in the oldest-old, given that age was top-coded in NHNAES IV and thus, we do not know their true chronological ages.

### Mortality

Mortality follow-up was based on linked data from records taken from the National Death Index through 2011, provided through the CDC [19]. Data on mortality status and length of follow-up (in person months) were available for all participants. Out of nine underlying causes of death and an “other category” that were provided in the linked data, eight were used to assess cause-specific mortality in our study—heart disease, cancer, chronic lower respiratory disease, cerebrovascular disease, diabetes, influenza or pneumonia, nephritis/nephrosis. Alzheimer’s disease was not considered in the cause-specific analysis due to the low number of cases. Accidents were not assessed due to the fact that many were probably not age-related and it is impossible to differentiate age versus non-age related accidental death.

### Phenotypic Age

Out of 42 possible clinical measures (details available on request) that were consistent across NHANES III and IV, variables were selected for inclusion in the Phenotypic Age measures (which we called Phenotypic Age) using a Cox Proportional Hazard Elastic Net model for mortality. Nine biomarkers (albumin, creatinine, glucose, (log) C-reactive protein, lymphocyte percent, mean cell volume, red blood cell distribution width, alkaline phosphatase, and white blood cell count) and chorological age were selected for inclusion based on ten-fold cross-validation.

Two Gompertz proportional hazard models were fit. One model, h_1_(t), was fit using all ten selected variables, and the other model, h_2_(t), was fit using only chronological age. “Phenotypic Age” used for predicting t-year mortality was calculated by asking what chronological age in h_2_(t) produced a hazard that was equal to the hazard for h_1_(t), estimated based on the individual’s observed levels of nine biomarkers and age. Thus, Phenotypic Age represents the expected age within the population that corresponds to a person’s estimated mortality risk (hazard at t years). Formula were expressed as following:

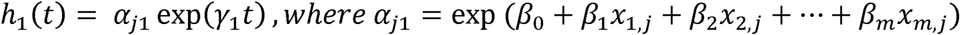

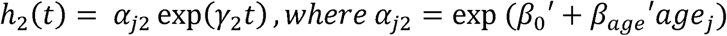

Suppose we want to get phenotypic age for prediction of mortality after 1 year,

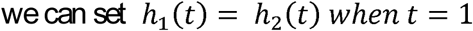

and then solve the following equation to get phenotypic age for subject j (*PhenoAge_j_*)

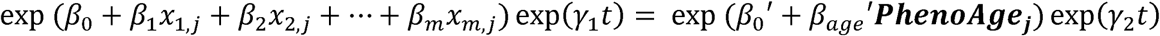

### BMI and disease count

BMI was calculated as weight in kilograms divided by height in meters squared. Normal BMI was defined as a value between 18.5 and 25.0. Chronic disease burden included a count of ten coexisting self-reported conditions: congestive heart failure, stroke, cancer (excluding skin), chronic bronchitis, emphysema, cataracts, arthritis, type 2 diabetes, hypertension, and myocardial infarction. Based on the disease counts, we created a variable with five categories—no disease, one disease, two diseases, three diseases, and four or more diseases.

### Statistical analyses

Age stratified ordinary least square (OLS) regression models were used to estimate the association between disease count and Phenotypic Age, within three age categories (20-39 y, 40-64 y, and 65-84 y). Based on these regression equations, we estimate the incremental increase in Phenotypic Age for each of the disease count categories in comparison to participants with no disease. Next, a parametric proportional hazard model (Gompertz distribution) was used to assess the association between Phenotypic Age and all-mortality, with adjustment for chronological age. To further evaluate robustness, age-stratified models and a model that excluded short-term mortality (within five years after baseline, to ensure that Phenotypic Age was not just capturing an end of life phenotype) were also run.

Next, participants were grouped into quintiles for the residual of Phenotypic Age (after regressing out chronological age), so that the highest quintile represented individuals who were most at-risk of death for their ages—i.e. those whose Phenotypic Age was the highest relative to their chronological age. We then plotted Kaplan-Meier Curves for persons in the highest 20% versus the lowest 20%. We also compared predicted median life expectancies at age 65 by sex and the five quintiles for Phenotypic Age, adjusted for age. Receiver Operating Characteristics (ROC) curves were used to compare ten-year mortality risk prediction of Phenotypic Age to individuals’ clinical chemistry measures and routine risk assessment tools.

Cause-specific mortality risk as a function of Phenotypic Age was assessed via Fine and Gray’s competing risk models [20]. To determine whether Phenotypic Age could differentiate risk in a healthy population, we conducted the all-cause mortality analysis again, restricting the sample to those with no reported disease and with normal BMI. Finally, participants aged 85+(oldest-old) were excluded from all prior analyses given that age was top-coded; therefore, to test mortality associations in this group, we used two parametric proportional hazard models (Gompertz distribution), one unadjusted and another with adjustment for disease counts, rather than chronological age (unknown).

All analyses were performed using R version 3.4.1 (2017-06-30) and STATA version 14.0 software (Stata Corporation, College Station, TX).

## Results

### Prevalence of disease

The basic characteristics of the study participants are shown in Table S1. Fig 1 presents the disease counts overall and by age category. Approximately two-thirds of the study participants were disease free at baseline, while about 22% had reported one chronic disease, 8.7% reported two diseases, about 3% reported three diseases, and only 1.9% reported at least four co-existing chronic diseases. As expected, the majority (86.6%) of young adults (ages 20-39 y) were free of disease, compared to about 60% of middle aged (40-64 y) and only a quarter of older adults. Additionally, nearly 7% of older adults had four or more chronic diseases, while only 1.25% of middle aged adults, and essentially no younger adults reported four or more disease diagnoses.

**Fig 1.**
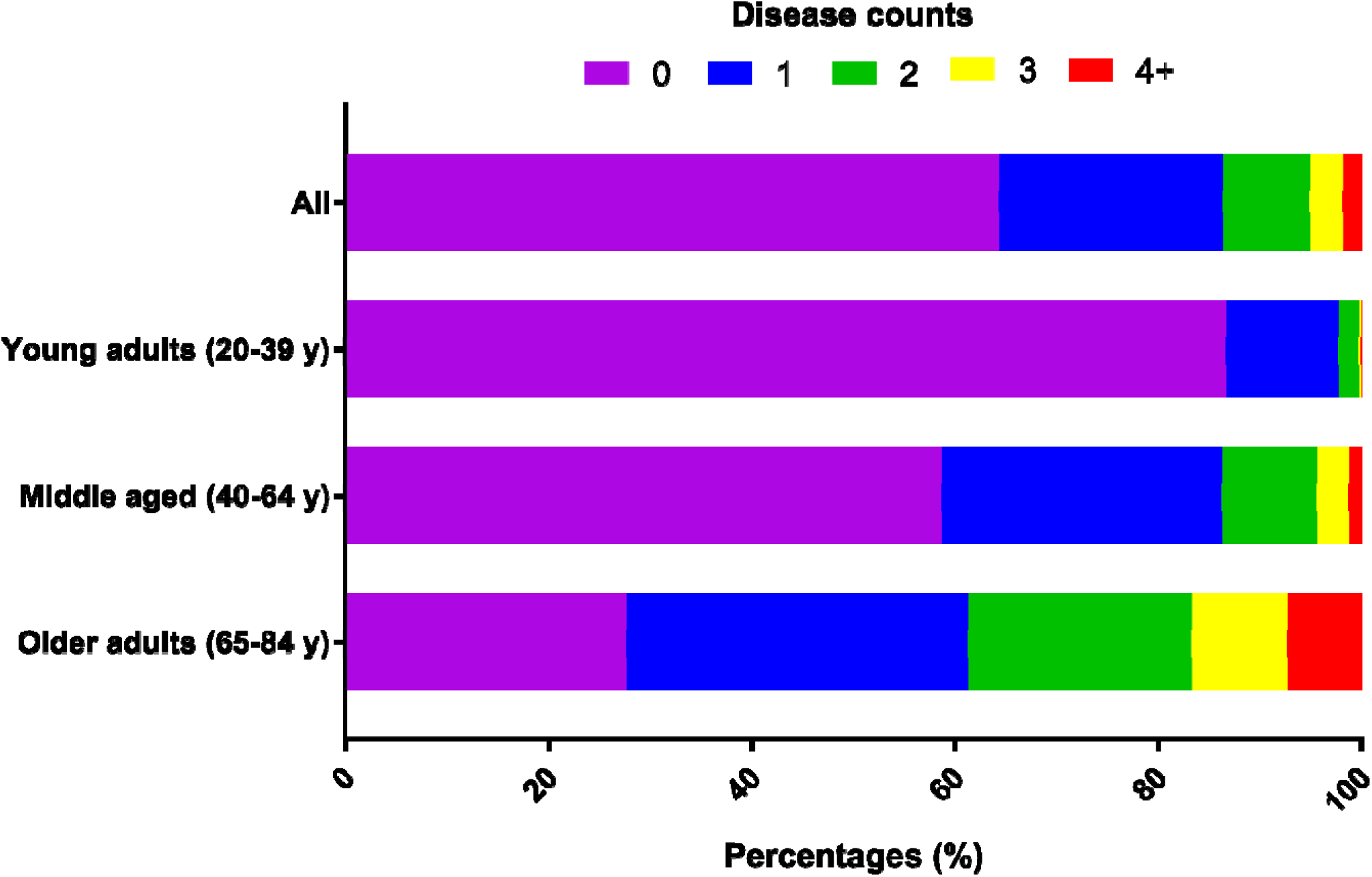
Frequency of disease counts overall and by age category.

### Phenotypic Age according to disease counts and age categories

Fig 2 shows the distribution of Phenotypic Age residual—residual of Phenotypic Age regressed on chronological age. A score of zero suggests a Phenotypic Age that was equivalent to one’s chronological age, whereas a positive value suggests a person looks older than expected, based on his/her phenotype for the nine clinical markers. Fig 3 shows predicted increases in Phenotypic Age for each disease count category, as compared to persons with no diagnosis of disease. Overall, participants with disease had older Phenotypic Ages compared to those without disease. For instance, among young adults, those with one disease were 0.2 years older phenotypically than disease-free persons, and those with two or three diseases were about 0.6 years older phenotypically. In middle aged adults, compared to those who were disease-free, those with one disease had Phenotypic Ages that were 0.2 years older, those with two diseases had Phenotypic Ages that were 0.3 years older, and those with three or more were on average 0.6-0.7 years older, phenotypically. Finally, for older adults, Phenotypic Age increased consistently as a function of disease, with those reporting one disease having Phenotypic Ages that were on average 0.1 years older than disease-free participants, those with two diseases having Phenotypic Ages 0.17 years older, those with three diseases having Phenotypic Ages 0.4 years older, and those with four or more diseases having Phenotypic Age that were 0.55 years older than disease-free older adults.

**Fig 2.**
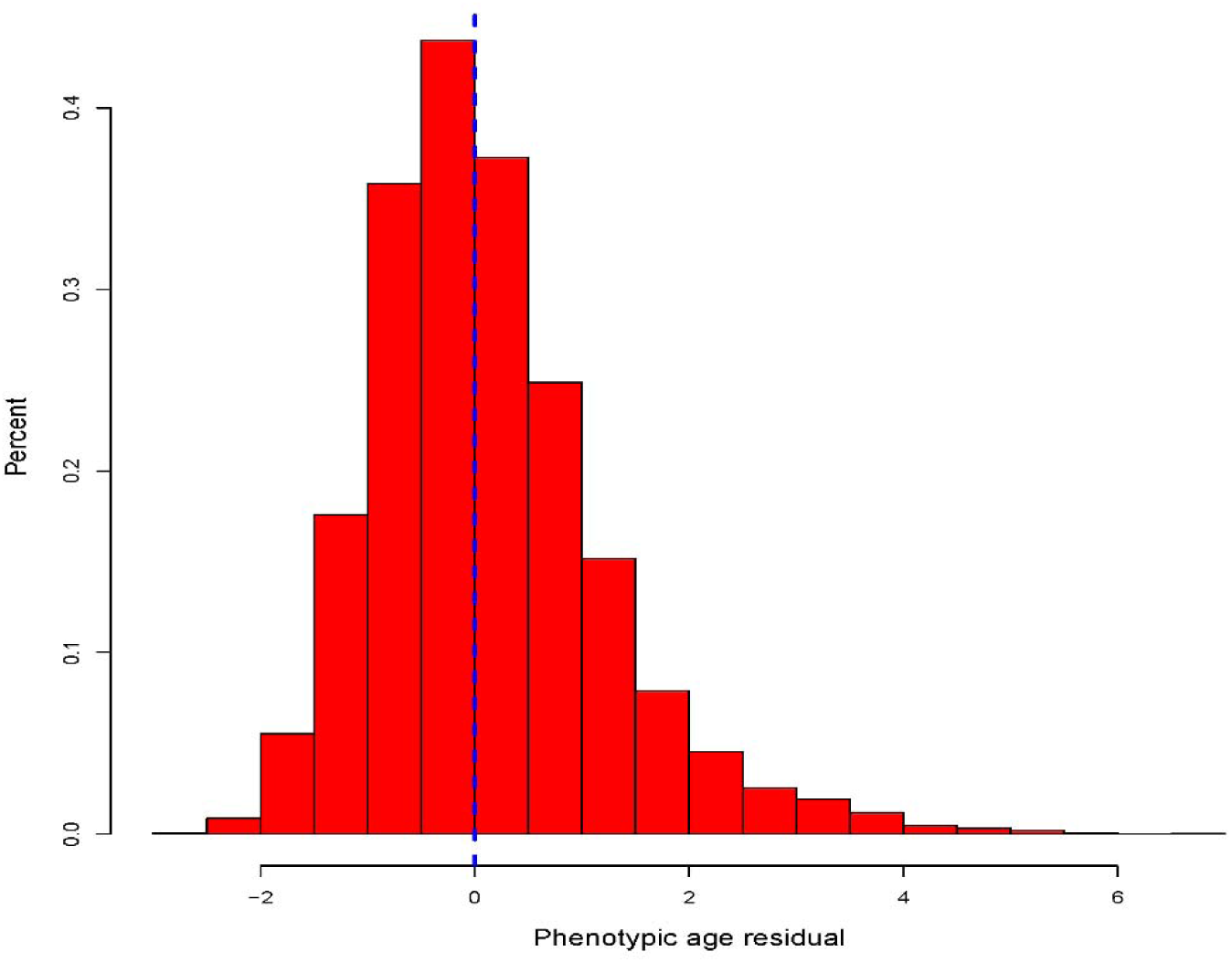
The distribution of Phenotypic Age residual. The x-axis depicts Phenotypic Age after taking the residual from a regression of Phenotypic Age regressed on chronological age. Mean and median Phenotypic Age is about zero, suggesting that on average participants had the same Phenotypic Age as their chronological age. A positive value represents being older than expected phenotypically, when taking into account one’s chronological age.

**Fig 3.**
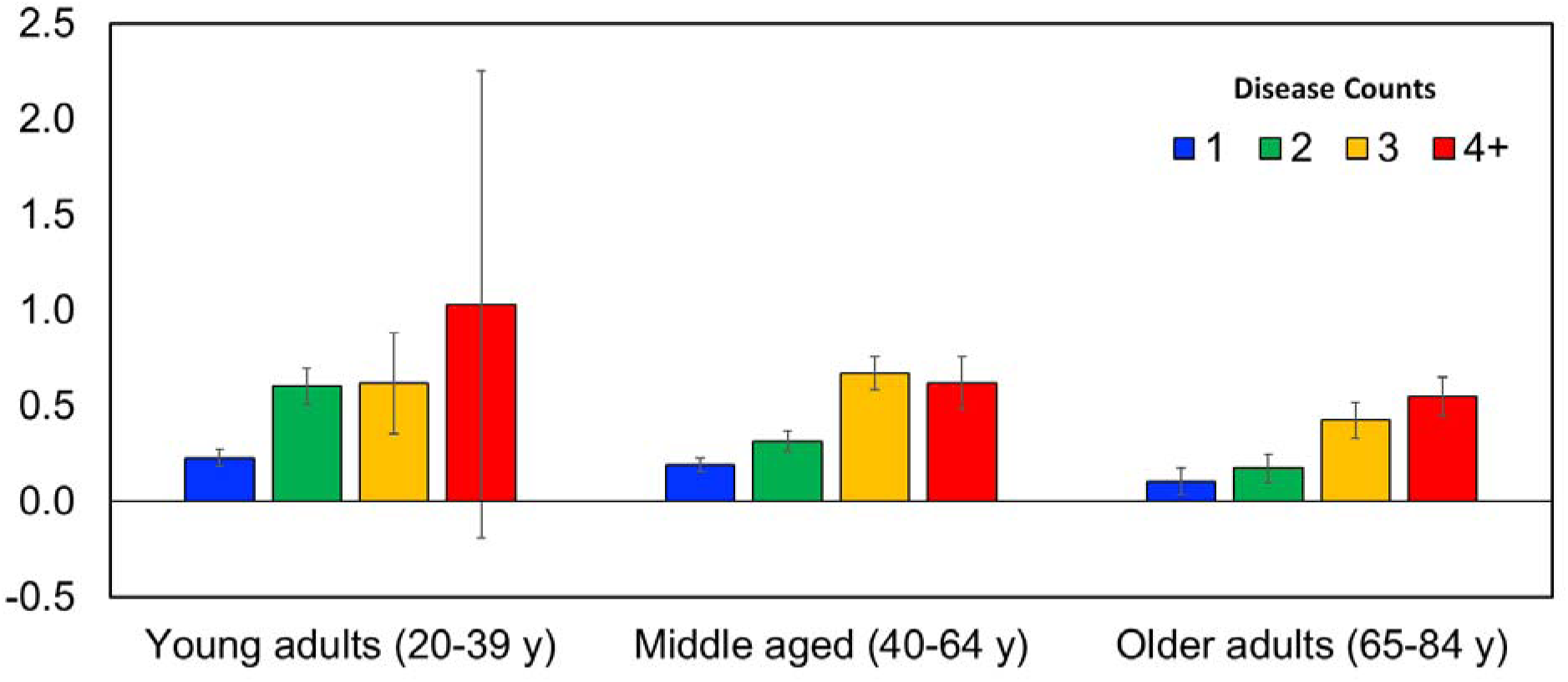
Predicted increases in the difference between Phenotypic Age and chronological age for each disease count by age category. The y-axis, depicts the increase in Phenotypic Age (after accounting for chronological age) as comparison to persons who were disease-free. The x-axis shows groups categorized based on chronological age and the number of diseases each participant had. For all age categories, we observed that Phenotypic Age was higher among persons who were diagnosed with more chronic diseases.

### Associations with all-cause mortality

Table 1 shows the associations between Phenotypic Age and all-cause mortality, based on proportional hazard models with Gompertz distribution. In the full sample, each one year increase in Phenotypic Age (after adjusting for chronological age) increased the risk of mortality by 9% (Hazard ratio [HR]=1.09, 95% confidence interval [CI]=1.08-1.10). When restricting the sample to participants who survived at least five years after baseline, we found consistent results, such that one year increase in Phenotypic Age was associated with an 8% increase in mortality risk. When examining mortality within age stratified groups, we found that Phenotypic Age was predictive in all age groups, such that each one year increase in Phenotypic Age was associated with a 14% increased mortality risk in young adults, 10% in middle-aged adults, and 8% in older adults. Finally, we found that on average, females were phenotypically younger than males (β=-1.34, P<2.2E-16); therefore, we compared sex-stratified model of all-cause mortality associations and found identical results for both sexes (females: HR=1.09, 95%CI=1.07-1.11; males: HR=1.09, 95%CI=1.07-1.11).

**Table 1.** Associations between Phenotypic Age and mortality.

|  |  | Phenotypic Age |  |  |  |
| --- | --- | --- | --- | --- | --- |
|  |  | No. of<br>deaths | Hazard Ratio<br>(95% CI) | z-score | P value |
| All-cause | Full sample | 873 | 1.09 (1.08-1.10) | 16.12 | <2.2E-16 |
|  | All-cause (5+ years survival) | 389 | 1.08 (1.06-1.10) | 8.28 | <2.2E-16 |
|  | Young adults (20-39 y) | 32 | 1.14 (1.09-1.18) | 6.66 | 2.7E-11 |
|  | Middle aged (40-64 y) | 247 | 1.10 (1.08-1.12) | 10.76 | <2.2E-16 |
|  | Older adults (65-84 y) | 592 | 1.08 (1.07-1.09) | 11.43 | <2.2E-16 |
| Disease-specific | Heart disease | 141 | 1.11 (1.08-1.14) | 8.26 | <2.2E-16 |
|  | Cancer | 227 | 1.07 (1.06-1.09) | 7.86 | 3.8E-15 |
|  | Chronic lower respiratory disease | 52 | 1.07 (1.04-1.11) | 4.42 | 9.9E-6 |
|  | Cerebrovascular disease | 56 | 1.04 (1.00-1.09) | 1.75 | 0.080 |
|  | Diabetes | 26 | 1.19 (1.12-1.26) | 6.03 | 1.6E-9 |
|  | Influenza or pneumonia | 24 | 1.12 (1.08-1.16) | 5.87 | 4.4E-9 |
|  | Nephritis/nephrosis | 15 | 1.20 (1.16-1.24) | 10.27 | <2.2E-16 |

CI, confidence interval. Results are based on Parametric Survival Models (Gompertz distribution). All models are adjusted for chronological age.

As shown in Fig 4, we found that those with the highest Phenotypic Ages, relative to their chronological ages had much steeper declines in survival over the approximately 12.5 years of follow-up. Interestingly, the high-risk groups appeared to have mortality rate that were similar, or in some cases higher than persons in the lowest risk group who were ten years older chronologically. For instance, among persons aged 50-64 y at baseline, about 25% of the high risk group had died after ten years of follow-up. Conversely among persons aged 65-74 y only about 20% of those in the low risk group had died after ten years of follow-up. For persons aged 65-74 y in the high risk group about half had died after ten years, compared to only about 67% of the low risk group who aged 75-85 y.

**Fig 4.**
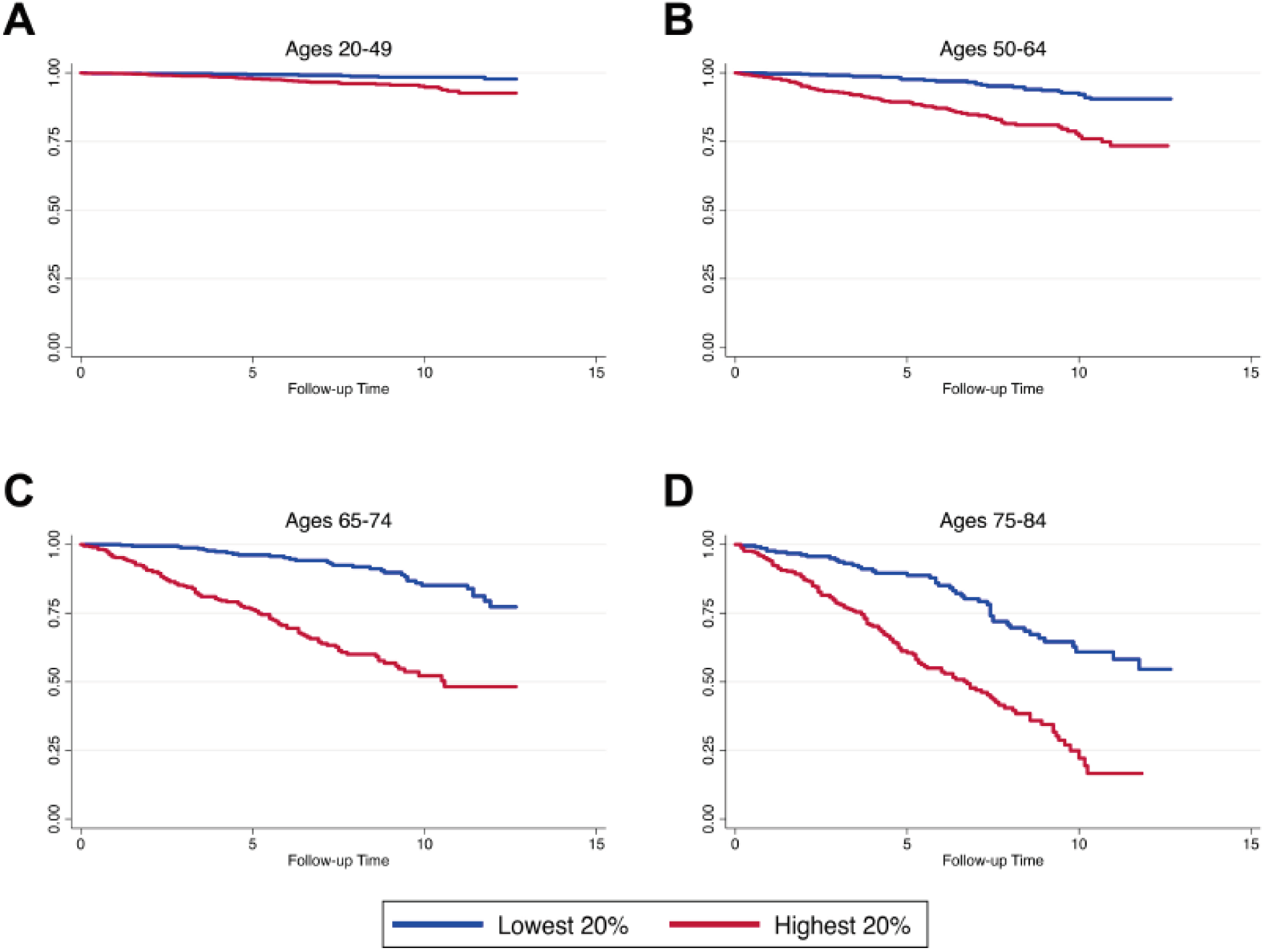
Kaplan-Meier Curves for persons in the highest 20% versus the lowest 20%. The y-axis indicates the survival rate; and x-axis indicates follow-up time (in month).

Fig 5 presents predicted median life expectancies at age 65 by sex and the five quintiles for Phenotypic Age. Results showed that 65-year-old females in the healthiest quintile had a predicted median life expectancy of about 87 y, while females in the unhealthiest quintile had a predicted life expectancy of just over 78 y. Similarly, 65-year-old males in the healthiest quintile had a predicted median life expectancy of about 84.5 y, while males in the unhealthiest quintile had a predictive life expectancy of just under 76 y.

**Fig 5.**
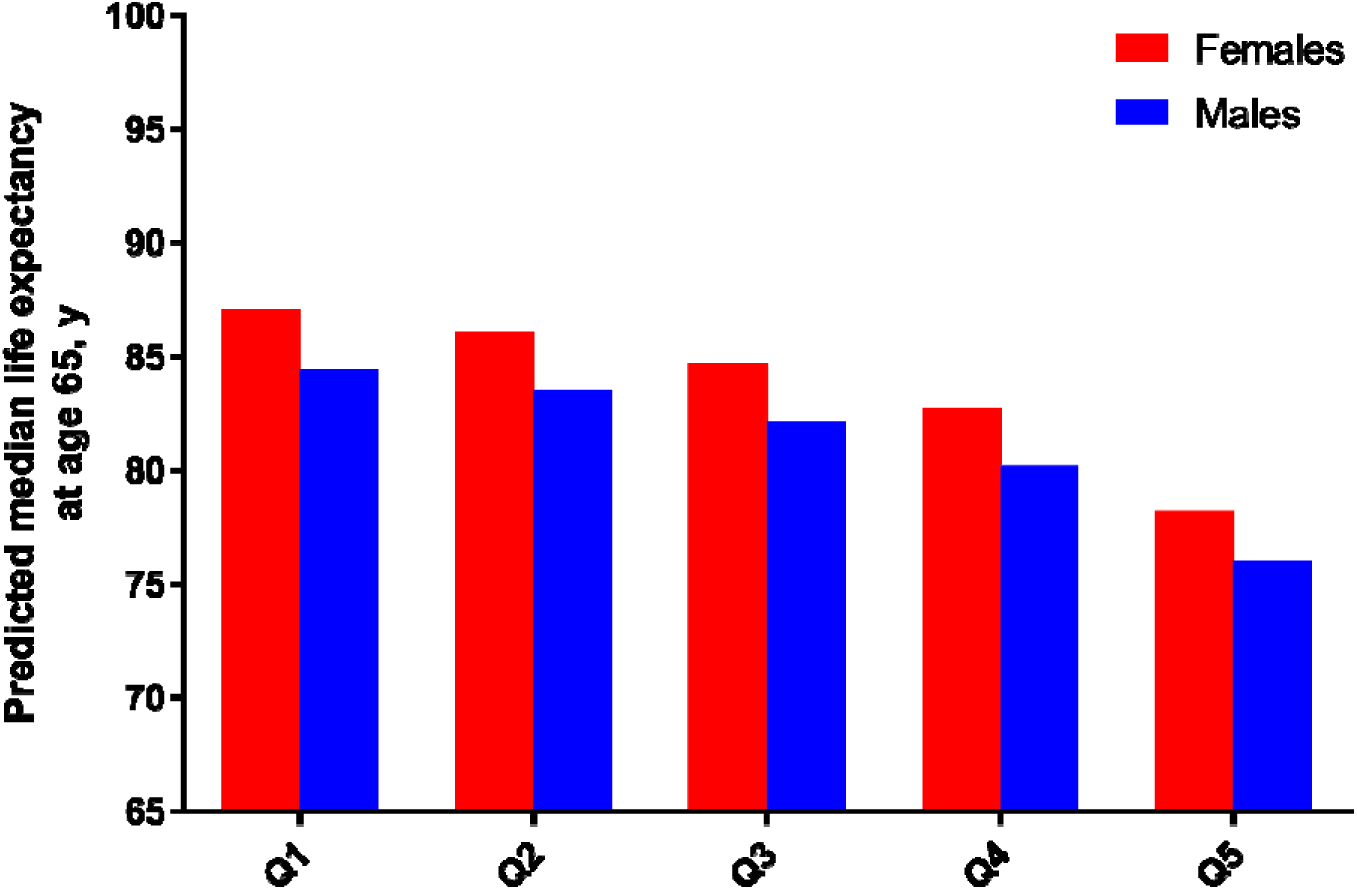
Predicted median life expectancies at age 65 by sex and the five quintiles for Phenotypic Age. Q1-Q5 indicates the five quintiles. Results are based on Parametric Survival Models (Gompertz distribution) that include quintiles of Phenotypic Age, chronological age, and sex. Estimates represent the predicted time in which 50% of the population is expected to have died, for each sex by quintile group, assuming a baseline age of 65.

ROC curves (Fig 6) revealed that Phenotypic Age significantly outperformed the other individual clinical chemistry measures, with an area under the curve (AUC) of 0.88. The next highest performing measures were chronological age with an AUC of 0.86, disease count with an AUC of 0.71, and serum creatinine with an AUC of 0.71. Four measures had AUCs between 0.60-0.69 (red cell distribution width, fasting glucose, systolic blood pressure, and albumin); five had AUCs between 0.50-0.59 (mean cell volume, lymphocyte percentage, C-reactive protein, alkaline phosphatase, and white blood cell count); and two had AUCs less than 0.50 (total cholesterol and BMI).

**Fig 6.**
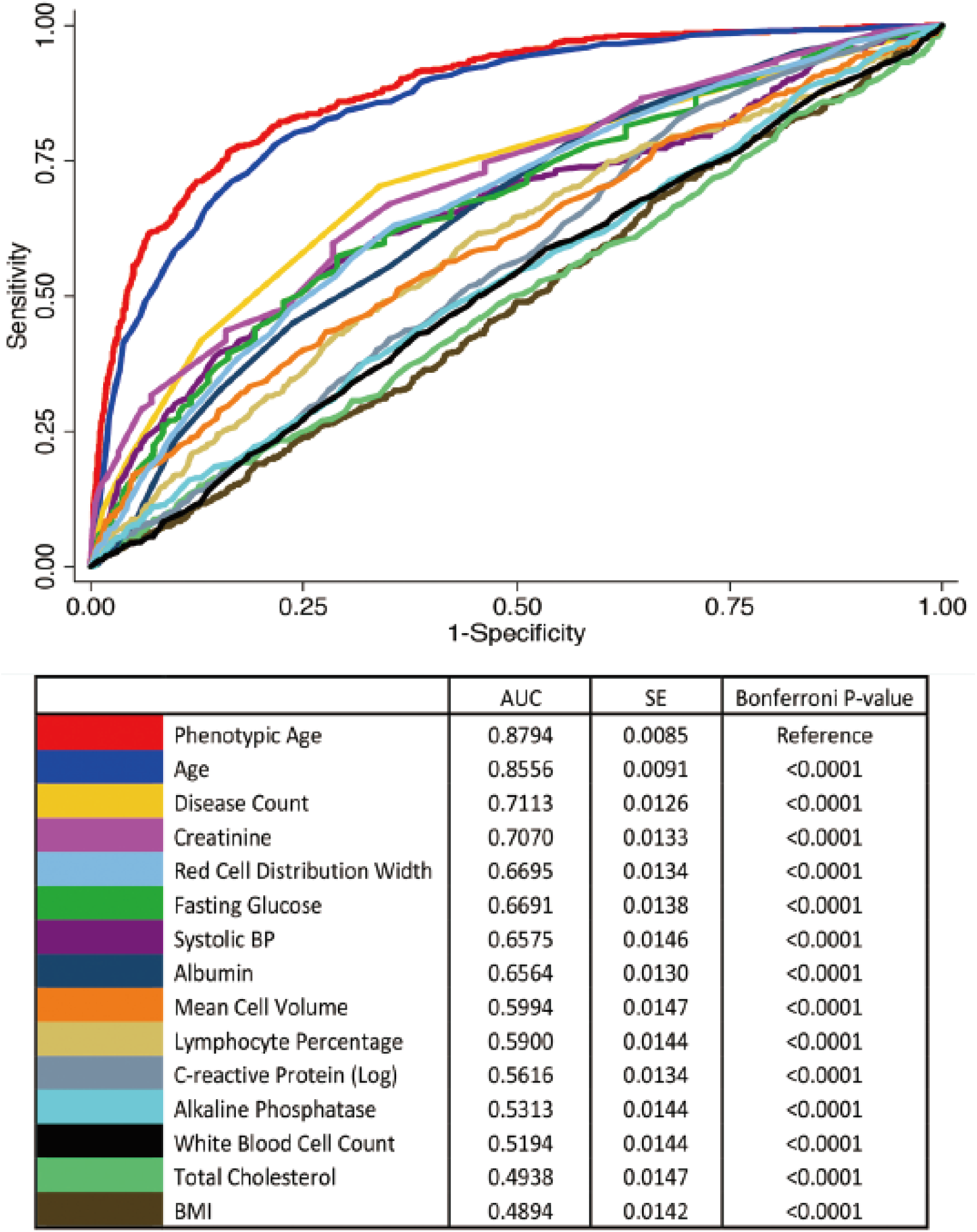
ROC curves for ten-year mortality. AUC, area under the curve; SE, standard error; BMI, body mass index.

### Associations with disease-specific mortality

As shown Table 1, Phenotypic Age was predictive of disease-specific mortality including heart disease, cancer, chronic lower respiratory disease, diabetes, influenza/pneumonia, and kidney disease. Conversely, a marginally significant association was found for cerebrovascular disease-specific mortality (HR=1.04, 95%CI=1.00-1.09).

### Associations with all-cause mortality in healthy participants

Given the need to identify aging measures that differentiate risk prior to disease pathogenesis, we tested all-cause mortality associations in participants free from disease and/or with normal BMI. As shown in Table 2, among those with normal BMI (n=3,243), we found that a one year increase in Phenotypic Age increased the risk of all-cause mortality by 9%, which is consistent with what was found for the full sample. Further, this effect remained when restricting our analytic sample to those with both normal BMI and no disease (n=1,906), such that a one year increase in Phenotypic Age was still associated with an 8% increase in all-cause mortality.

**Table 2.** Phenotypic Age and all-cause mortality in participants with normal BMI or normal BMI and disease free.

|  | Phenotypic Age |  |  |
| --- | --- | --- | --- |
|  | Hazard Ratio | z-score | P value |
|  | (95% CI) |  |  |
| With normal BMI | 1.09 (1.06-1.11) | 6.90 | 5.2E-12 |
| With normal BMI and disease free | 1.08 (1.03-1.14) | 2.94 | 0.003 |

BMI, body mass index; CI, confidence interval. Results are based on Parametric Survival Models (Gompertz distribution). All models are adjusted for chronological age. Normal BMI was defined as a value between 18.5 and 25.0.

### Associations with all-cause mortality in the oldest-old

Table 3 provides the mortality associations in the oldest-old. We found that regardless of adjustment Phenotypic Age was associated with mortality in this subpopulation, although to a lesser degree than the full population (Unadjusted Model: HR=1.05, 95%CI=1.02-1.09; Disease Adjusted Model: HR=1.04, 95%CI=1.01-1.18).

**Table 3.** Phenotypic Age and all-cause mortality in the oldest-old (aged 85+)

|  | Phenotypic Age |  |  |
| --- | --- | --- | --- |
|  | Hazard Ratio | z-score | P value |
|  | (95% CI) |  |  |
| Model 1: Without disease count adjustment | 1.05 (1.02-1.09) | 3.28 | 0.001 |
| Model 2: With disease count adjustment | 1.04 (1.01-1.18) | 2.26 | 0.024 |

CI, confidence interval. Results are based on Parametric Survival Models (Gompertz distribution). Models were not adjusted for chronological age, given that it was top-coded at age 85 in NHANES IV.

## Discussion

In a national-representative US adult population, we showed that our new measure of aging—Phenotypic Age—was highly predictive of remaining life expectancy using an independent validation sample. Overall, we found that the mortality prediction of this measure is valid across different stratifications by age, health, sex, and cause of death. For instance, it is strongly associated with all-cause mortality in multiple age groups, including young adults, middle-aged, and older adults. Moreover, the effect sizes seem to decrease with age, which may suggest that in younger groups, when the risk of death is low, variations in physiological status—as captured by Phenotypic Age—may play a bigger role in who lives longer. Conversely, in older adults, for whom the risk of death increases, mortality may be more stochastic. Nevertheless, we were able to determine that this measure was not just capturing an end-of-life or critically ill status, given that it remained predictive of mortality after excluding participants who had not survived for at least five years after baseline.

The finding that Phenotypic Age was predictive of mortality among both healthy and unhealthy populations is novel. Many of the measures of aging, such as those based on deficit accumulation [21, 22], include measures of morbidity in their construction, and thus it is impossible to disentangle aging and disease, or determine the usefulness of such measures in healthy populations. Belsky et al. evaluated aging measures, including our previous measure, in a cohort study of young adults, who were mostly disease-free [5, 16]. However, the outcomes available were mostly restricted to functional assessments, which may mean something different in younger adults than they do in older populations. Conversely, in this study, we were able to show that Phenotypic Age was predictive of all-cause mortality among disease-free, healthy older adults. This suggests that Phenotypic Age is not simply a measure of disease or morbidity and instead may be a marker that tracks the effect of aging before diseases become clinically evident. This suggests that in a clinical setting, Phenotypic Age could be used to stratify risk among persons who otherwise “appear” healthy.

As expected of an aging biomarker, Phenotypic Age also tracks multimorbidity. We observed a strong association between the number of diseases a person reported being diagnosed with and his/her Phenotypic Age, relative to his/her chronological age. Despite relatively small sample sizes, in general, Phenotypic Age appeared increased as a function of disease count, suggesting that among persons of the same age, the more coexisting diseases a person has, the more phenotypically older he/she appears—based on clinical biomarkers. Nevertheless, Phenotypic Age predicted risk of death significantly better than disease count, suggesting that it is capturing information beyond a person’s number of co-existing conditions. Further, Phenotypic Age was also associated with mortality in the oldest-old, which is a highly disease prevalence population, and this association remained even after adjusting for disease count. While we did not have follow-up data on disease incidence, one explanation that should be examined further is whether higher phenotypic age is predictive of disease accumulation. For instance, among persons with one disease, an important question is whether Phenotypic Age predicts who will develop a second comorbid condition.

The efficacy of Phenotypic Age to assess mortality risk in the general population, as well as multiple subpopulations that are heterogeneous in age and health status, provides strong evidence of its suitability for applications in both the clinical setting and research in the biology of aging. For instance, the generalizability of Phenotypic Age to assess risk of various aging outcomes, may facilitate identification of at-risk individuals for a number of distinct conditions. Phenotypic Age may also be a useful marker for evaluation of interventions—particularly those concerned with prevention by delaying disease pathogenesis [15, 23–25]. Aging changes are hypothesized to begin as early as conception [26]—preceding disease—thus interventions to slow aging will be most effective for reducing disease incidence if started early in the life course prior to significant accumulations in aging-related damage. Our findings suggest that Phenotypic Age is in-line with the Geroscience paradigm, which stipulates that “aging is the greatest risk factor for a majority of chronic diseases driving both morbidity and mortality” [27, 28]. Therefore, measures such as Phenotypic Age that capture pre-clinical aging as well as future morbidity/mortality risk could facilitate evaluation of intervention efficacy, while avoiding the need for decades of follow-up [23]. While research to develop interventions that target the aging process is ongoing, our paper provides a potential end-point for which they can be evaluated. Further, this metric may also shed light on factors that alter the pace of aging, facilitating investigation into potential biological mechanism and environmental stressors.

In conclusion, our study shows that Phenotypic Age, a novel clinically based measure of aging, is a consistently strong predictor of all-cause mortality in a nationally-representative population. Importantly, the measure is robust to population characteristics—it is a reliable mortality predictor regardless of the age or health status of the population being assessed. Further, this measure captures both all-cause and disease-specific mortality, and is also strongly associated with the number of comorbid conditions. In moving forward, measures such as this may facilitate both secondary and tertiary prevention, via identification of at-risk groups and evaluation of intervention efficacy. Phenotypic Age has the potential to also serve as an outcome variable in basic research, with the goal of identifying molecular mechanisms that regulate the rate of aging.

## Acknowledgments

This study was funded by a grant from US NIH/NIA (4R00AG052604-02).

